# Improving retention time of Indocyanine Green for *in vivo* two-photon microscopy using liposomal encapsulation

**DOI:** 10.1101/2025.03.20.644063

**Authors:** Alankrit Tomar, Noah Stern, Tyrone Porter, Andrew K. Dunn

**Author notes:** co-first author.

## Abstract

We propose a technique for improving the retention time of Indocyanine Green (ICG), an FDA-approved, widely available, near infrared (NIR) probe for *in vivo* imaging of the cerebrovascular structure in rodents. We present a synthesis process for liposomal nanoparticles that, when used to encapsulate ICG, significantly increase the circulation time of the vascular label. Further, we conduct *in vivo* imaging experiments with unencapsuated (free) ICG and liposomal ICG (L-ICG) and compare the retention of ICG in the vascular network. Our work tackles one of the most fundamental limitations in using ICG i.e. rapid clearance, for the purpose of two-photon microscopy.

## 1 Introduction

Two-photon fluorescence microscopy (TPM) is widely used to visualize the cerebrovascular structure of the rodent brain and other tissues at high spatial resolution and at extended depths.^1, 2^ The imaging procedure for blood vessels involves an intravenous injection of a fluorophore that labels the blood plasma in the blood vessels. The fluorescent vascular label can then be excited via a two-photon excitation process. Common vascular labels for two-or three-photon microscopy include dextran-conjugated Texas Red,^3, 4^ Fluorescein isothiocyanate-conjugated dextran (FITC-dextran),^5^ and Alexa Fluor dyes.^6–8^ In the past, deep and extended two-photon imaging of the vascular network has been illustrated using these fluorescent dyes.

However, these widely used probes, while effective, are often expensive. A single intravenous injection in a mouse for dextran-conjugated Texas Red costs around $50 and in the range of $400-500 for the Alexa Fluor dyes, depending on the specific variant. On the other hand, Indocyanine Green (ICG) is a relatively inexpensive alternative that is also FDA-approved. Compared to the aforementioned vascular dyes, a single intravenous injection of ICG in a mouse costs just 80 cents. In addition, due to its wide utility for clinical application, extensive research is available on the efficacy, safety, and use cases for ICG. This serves as motivation for adoption of ICG as a vascular label for TPM.

Despite these advantages, there are two key limitations of ICG as a vascular label specifically for TPM. The first is that the peak two-photon excitation wavelength for ICG overlaps with the one-photon water absorption window around 1450 nm, making traditional two-photon excitation unfeasible.^9^ The second limitation is the rapid clearance of ICG from the vasculature. Miller et al. have previously noted that for two-photon microscopy in mice, the retention time for ICG is in the range of 20-25 minutes.^10^ The low retention time does not allow for longer imaging experiments. In this work, we propose that encapsulation of ICG in liposomal nanoparticles can help increase the retention or circulation time and enable improved two-photon fluorescence imaging.

Lipid-based nanoparticles, specifically liposomes, are one of the most heavily researched and commercialized nanoparticle systems.^11^ They have seen widespread use, approval, and licensing for everything from their earliest use to encapsulate doxorubicin for cancer therapy^12^ to their more recent use as carriers for the mRNA-based COVID-19 vaccines.^13^ Liposomal nanoparticles are inexpensive, biocompatible, biodegradable, easy to prepare, relatively stable, and highly scalable. Previous success has pushed a vast majority of lipid-based nanoparticle research to continue its focus on drug delivery, but increased circulation time and lower toxicity also suggests that liposomes may also be excellent carriers for exogenous contrast agents like ICG.^14^

Liposomes are an ideal candidate to prolong the circulation of ICG as they do not significantly shift or dampen the absorbance or fluorescence properties of the ICG.^15, 16^ In fact, several groups are currently exploring the use of liposomal ICG (L-ICG) in various forms and compositions for a variety of imaging modalities including fluorescent imaging,^17–21^ multispectral optoacoustic tomography,^15, 22^ photoacoustic imaging,^23–26^ and photothermal therapy^27–29^ Applications span several organ systems including the eyes^30^ and brain^31^ as well as multiple use cases including imaging the vasculature system and monitoring tumor progression. Separately Kraft and Ho^20^ and Gao et al.,^17^ show that encapsulation within liposomes actually improves the stability and fluorescence intensity of ICG for *in vivo* applications.^17, 20^ In the latter case, L-ICG show a 38.7-fold increase in near infrared fluorescence intensity as compared to free ICG likely due to the dense packing of individual ICG molecules in the liposomal shell.

In spite of these advantages, capitalizing on the clearance time benefits of liposomal encapsulation for two-photon microscopy of microvasculature in the brain has not been explored. Here, we detail a procedure for the formation of ICG-encapsulated liposomes along with relevant characterization. We perform independent *in vivo* imaging of the cortical vascular network labeled with free ICG and liposomal ICG (L-ICG). Finally, we undertake a quantitative comparison between the clearance time of free ICG and L-ICG.

## 2 Methods

### 2.1 Synthesis of liposomal nanoparticles

Encapsulation of ICG within liposomes consists of three major steps: thin-film hydration, extrusion, and dialysis. Briefly, a thin-film of lipids DSPC, cholesterol, and DSPE-PEG2k is formed in a defined ratio along the walls of a glass vial under vacuum. Next, the film is hydrated with a high concentration solution of ICG and is gently heated and agitated to encourage nanoparticle formation. Once sufficiently hydrated, the resulting nanoparticle solution is extruded through 100 nm membranes to cut them down to size and increase uniformity. After extrusion, the liposomal solution is dialyzed with ultrapure water to remove any unencapsulated ICG. For *in vivo* applications, the nanoparticle solution is passed through sterile membranes and centrifuged to bring them to the desired concentration (1 mg/ml) for injection. A schematic of the entire process is shown in 1. The synthesis process is described in further detail in the supplemental material section.

**Fig 1.**
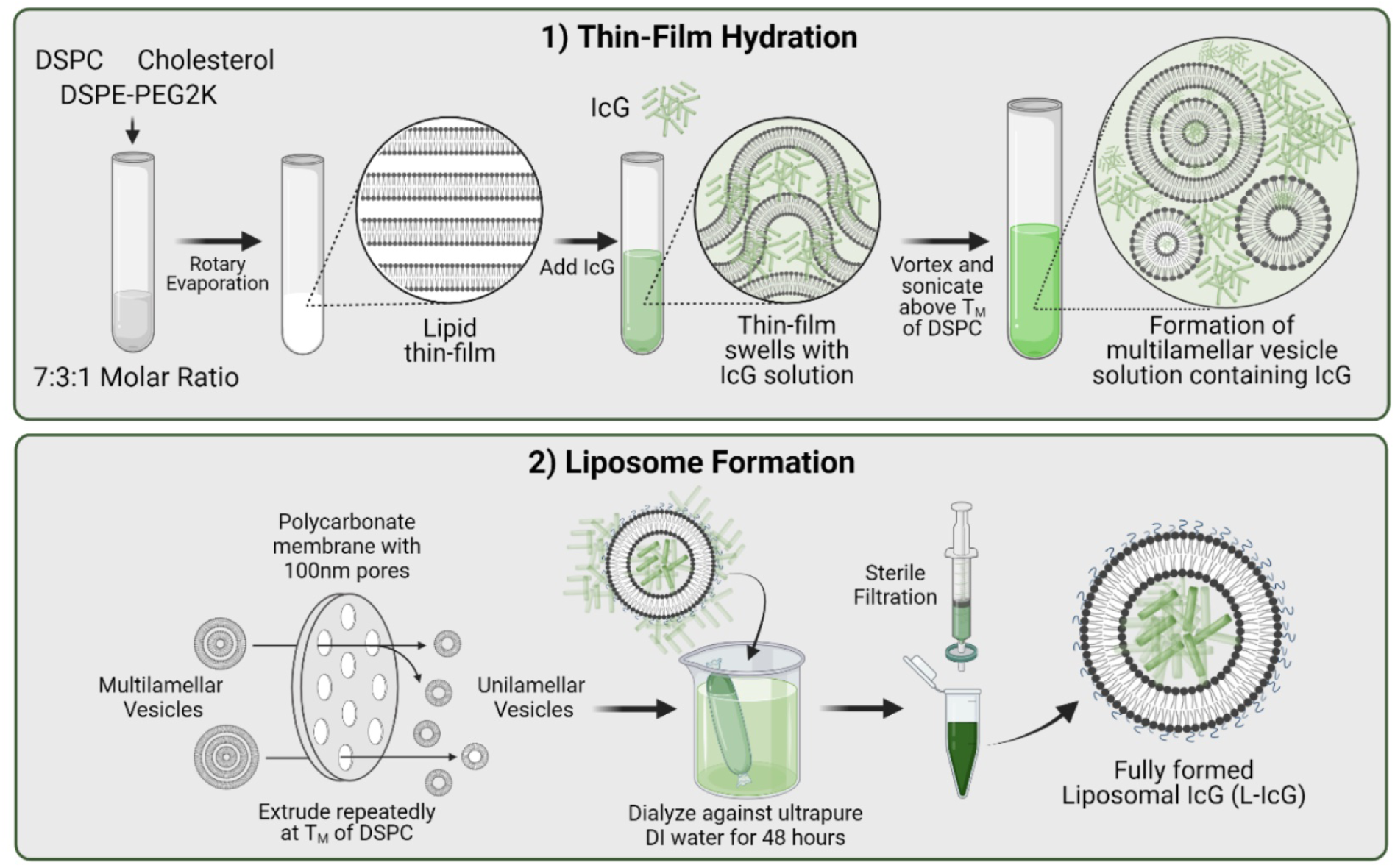
Schematic depicting the two-step process for synthesis of liposomal ICG particles. Phase 1 encompasses thin film hydration. In this phase, lipids are added at a defined concentration, evaporated to form a thin film and hydrated with a solution containing ICG. The resulting solution containing multilamellar vesicles with ICG moves to Phase 2 where liposomes are formed. In this phase, particles are extruded down to the proper size forming liposomes, dialyzed to remove any excess ICG, and sterile filtered to remove any contaminants and larger aggregates that still remain. For *in vivo* experimentation, an additional step of centrifugation and resuspension to the desired final concentration in sterile saline or PBS is performed. Refer to the supplemental material for a detailed description of the synthesis process. Schematic created with BioRender.

### 2.2 Characterization of liposomal nanoparticle

Size, zeta potential, particle concentration, encapsulation efficiency, and single photon fluorescence intensity are the main areas of characterization. Size, zeta potential, and particle concentration are obtained through analysis with a Malvern Zetasizer and NanoSight. Encapsulation efficiency is calculated as a ratio of the initial amount of ICG used during thin-film hydration to the final amount of ICG encapsulated within the nanoparticle. Concentrations of ICG are determined after mixing in a 1:1 ratio with ethanol and comparing to a standard curve. Single photon fluorescence intensity is measured using a Shimadzu spectrofluorophotometer. The characterization results and details are described in the supplemental material section.

### 2.3 Ultrafast laser sources

The laser source setup consists of a non-collinear optical parametric amplifier (NOPA, Spectra Physics) which was pumped by a 1 MHz repetition rate laser (Spirit 1030-70, Spectra-Physics) at 35 W. The NOPA offered independent tunability of the wavelength across a wide range of wavelengths, but for the purpose of the experiments described in this work, the output wavelength was set at 1300 nm.

### 2.4 Multiphoton Microscope

Imaging was performed using a custom home-built two-photon microscope.^3, 9^ A schematic of the microscope layout is shown in 2. Longpass filters (LPF 1: FEL0850 (Thorlabs), LPF 2: FELH1150 (Thorlabs), LPF 3: FF01-937 (Thorlabs)) were used to remove any unwanted residual wavelengths in the NOPA output. A combination of half-wave plate (HWP) and polarizing beam splitter (PBS) are used to modulate the power of the excitation beams during the experiment. The excitation beam was raster scanned during the experimental procedure with two galvanometric scan mirrors. A scan lens (f=50 mm, SL50-3P, Thorlabs) and a Plössl tube lens were used in conjunction (f=200 mm, 2x AC508-400-C, Thorlabs) to expand the beam to fill the back aperture of a 25x objective (XLPLN25XSVMP2, 1.0 NA, Olympus). The objective focuses the excitation light onto the sample leading to emission of fluorescence signal. The emitted photons were reflected towards the detection path by a dichroic mirror (FF980-DI01-T1, Semrock) wherein they passed through a set of emission filters (893/209, Semrock; 855/210, Semrock) before getting collected by photomultiplier tubes (H10770PB-50, Hamamatsu). The control of the stages, scanners, and image acquisition was carried out by a custom-made software in LabView and the image analysis was undertaken using Fiji.

**Fig 2.**
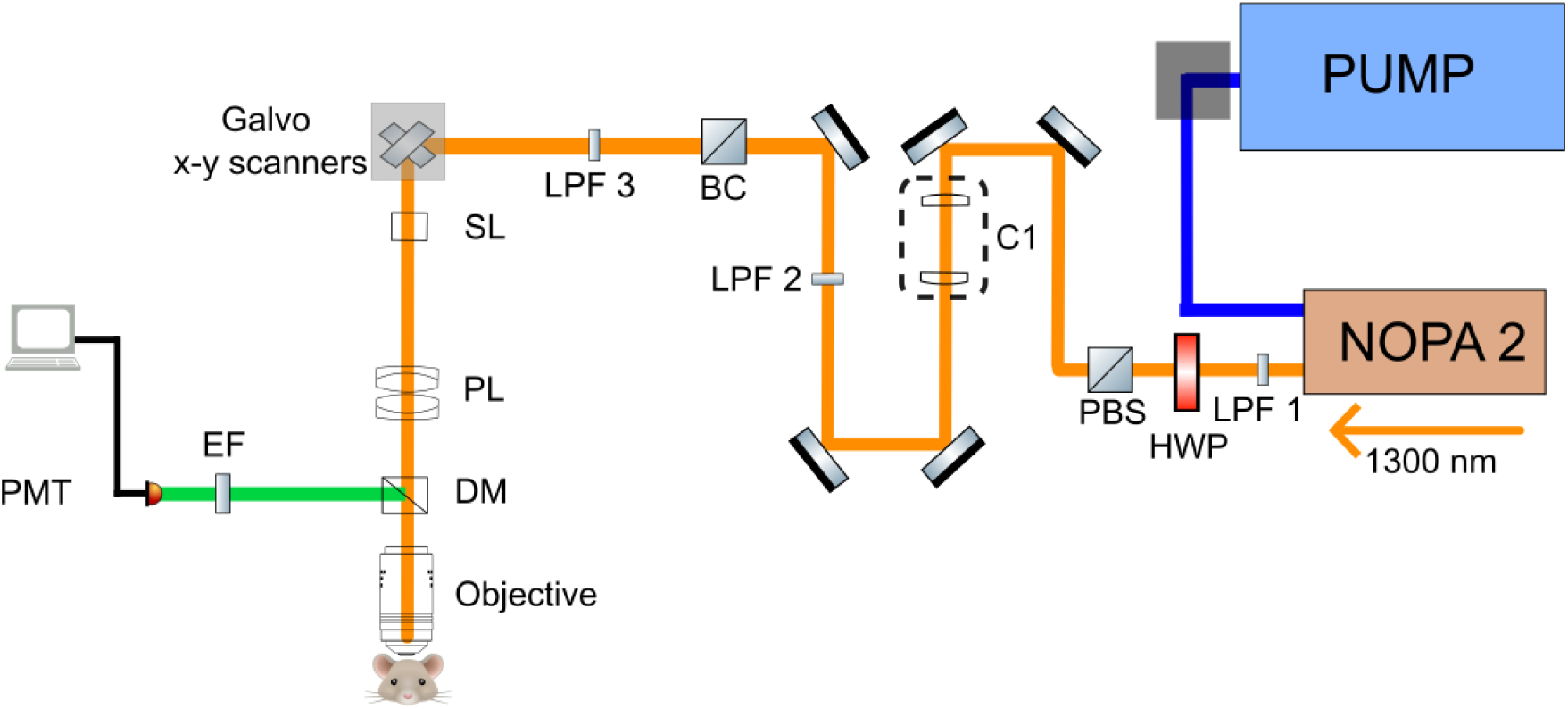
Layout of laser and imaging setup for two-photon microscopy. NOPA: noncollinear parametric amplifier, LPF: long pass filter, HWP: half wave plate, PBS: polarizing beam splitter, C: collimator, BC: beam combiner, SL: scan lens, TL: Tube lens, DM: Dichroic Mirror, EF: emission filters, PMT: photomultiplier tube

### 2.5 Animal Protocols

Institutional Animal Care and Use Committee at the University of Texas at Austin approved all the animal procedures. Isoflurane was used as our anesthetic agent (1.5-2%) for all surgical procedures. The cranial window site was chosen to the right of the sagittal suture between the coronal and lambdoidal sutures. A circular flap, approximately 5 mm in diameter, was drilled over which the cranial window was positioned. The mouse’s temperature was maintained at 37.6 ° C during the craniotomy procedure. Imaging procedures were performed after allowing the mice a rest of 2 weeks.

## 3 Results

### 3.1 In vivo procedures and signal variation with time

The objective of the *in vivo* studies was to compare the clearance of free ICG and L-ICG from the rodent vascular network as a function of time. A total of five *in vivo* imaging procedures were performed using L-ICG, the results for one of which are discussed below. On the other hand, two *in vivo* imaging procedures were undertaken using free ICG. The supplemental material section shows the *in vivo* imaging results obtained with L-ICG from two other mice. In this work, the term *clearance time* refers to the time after injection until reasonable vascular images can be obtained. Although this is a subjective definition, accurately quantifying clearance time is not crucial; it serves merely as a metric to compare ICG retention in the body.

The L-ICG compound was prepared to a concentration of 1 mg/ml. A solution of free ICG with a similar concentration was also prepared. The blood plasma of the vascular network in the cortex was labelled through a 200 µL retro-orbital injection of either the L- or free ICG solution. Imaging was performed with an excitation wavelength of 1300 nm, which is a suitable wavelength for excitation of ICG-based compounds. The Discussion section provides more information on the choice of wavelength used. During imaging, the average power incident on the surface was increased with depth but did not exceed 40 mW.

After injection, the first few minutes were spent setting up the mouse under the multiphoton microscope and finding suitable regions for imaging at different depths. Imaging of the vascular network began 15 minutes after the injection and continued at regular 15-minute intervals up to a total of 75 minutes. The deepest depth investigated was 600 *µ*m for all mice that were imaged.

Figures 3(a) and 4(a) show the imaging results from the experiments with free ICG and L-ICG, respectively. Note that within the same figure, the images are displayed on the same histogram scale. The clearance of the dye over time can be qualitatively inspected from the images but there is value in quantitatively analyzing the results. For the quantitative analysis, fluorescence intensity of vessels was tracked over time. Specifically, the mean fluorescence intensity for each vessel was determined by averaging the five highest pixel intensities from the line profile of the vessel. Additionally, when it was possible to track and quantify multiple vessels, 2-3 similarly sized vessels were selected from each image, and their mean fluorescence intensities were averaged to obtain a value representing the overall mean fluorescence intensity of the vessels at a given depth. The overall mean fluorescence intensity at each depth was normalized relative to the value at the initial time point, i.e., 15 minutes. For a certain depth, this will be referred to as the normalized fluorescence intensity. When possible, the signal to background ratios (SBR) for the vessels were also investigated. The SBR is quantified as (*µ*_sig_)/*µ*_bg_ where *µ*_sig_ denotes the mean fluorescence intensity (as defined above) and *µ*_bg_, mean background intensity, is found by averaging intensity values of 10 pixels on either end of the line profiles. The overall SBR (denoted by SBR in the figures) for a given depth at a specific time point was calculated by averaging the SBRs of all vessels analyzed at that depth. Note that an SBR of 1 denotes that the signal cannot be differentiated from the background.

**Fig 3.**
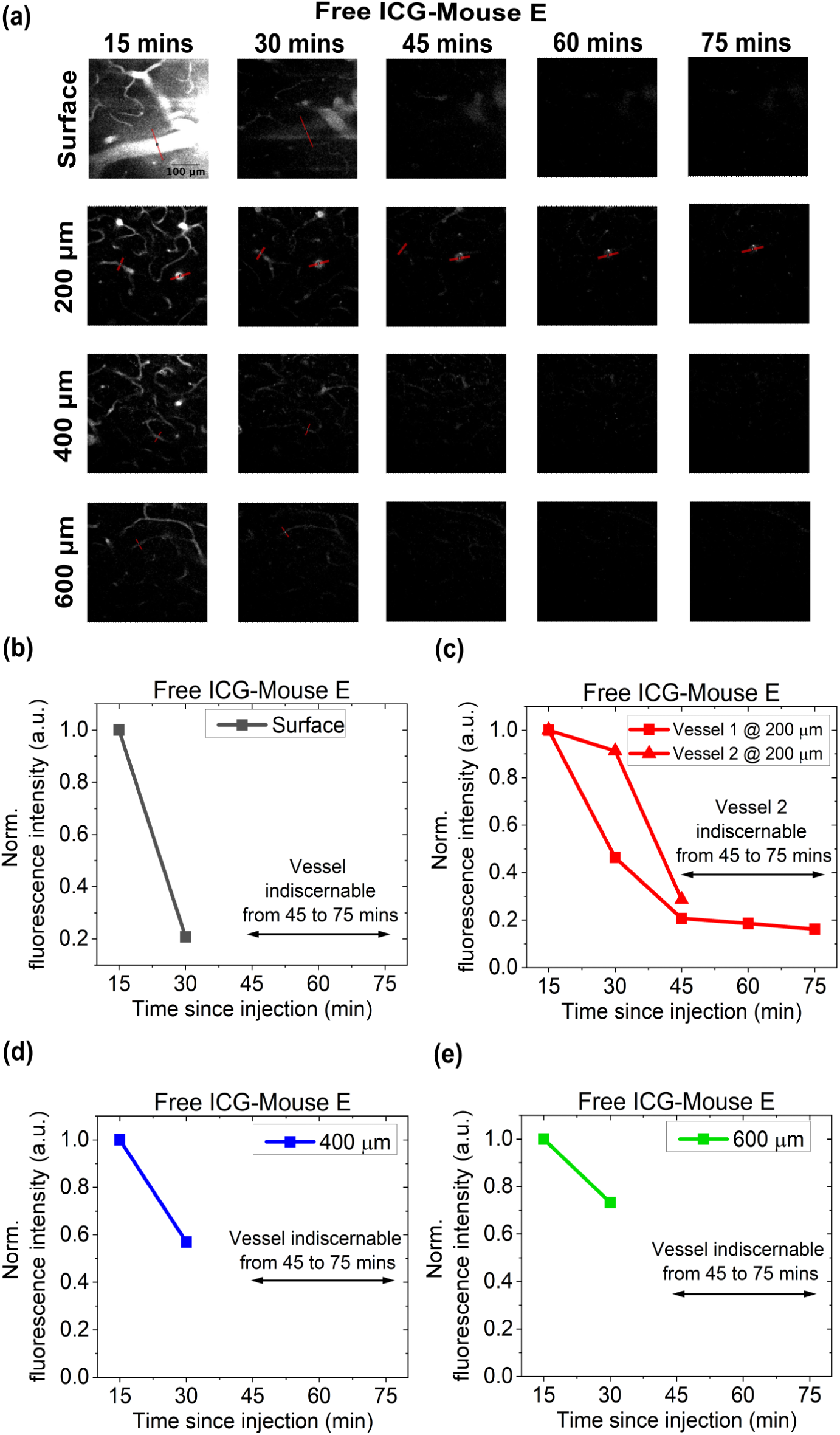
(a) *In vivo* imaging of blood vessels at different depths and time points for Mouse E using free ICG. All images are displayed on the same histogram display range. Scale bar represents 100 *µ*m. Variation of overall mean fluorescence intensity for vessels at surface (b), 200 *µ*m (c), 400 *µ*m (d), 600 *µ*m (e). The mean vessel fluorescence intensity at each time point is normalized to the value at the 15-minute time point.

**Fig 4.**
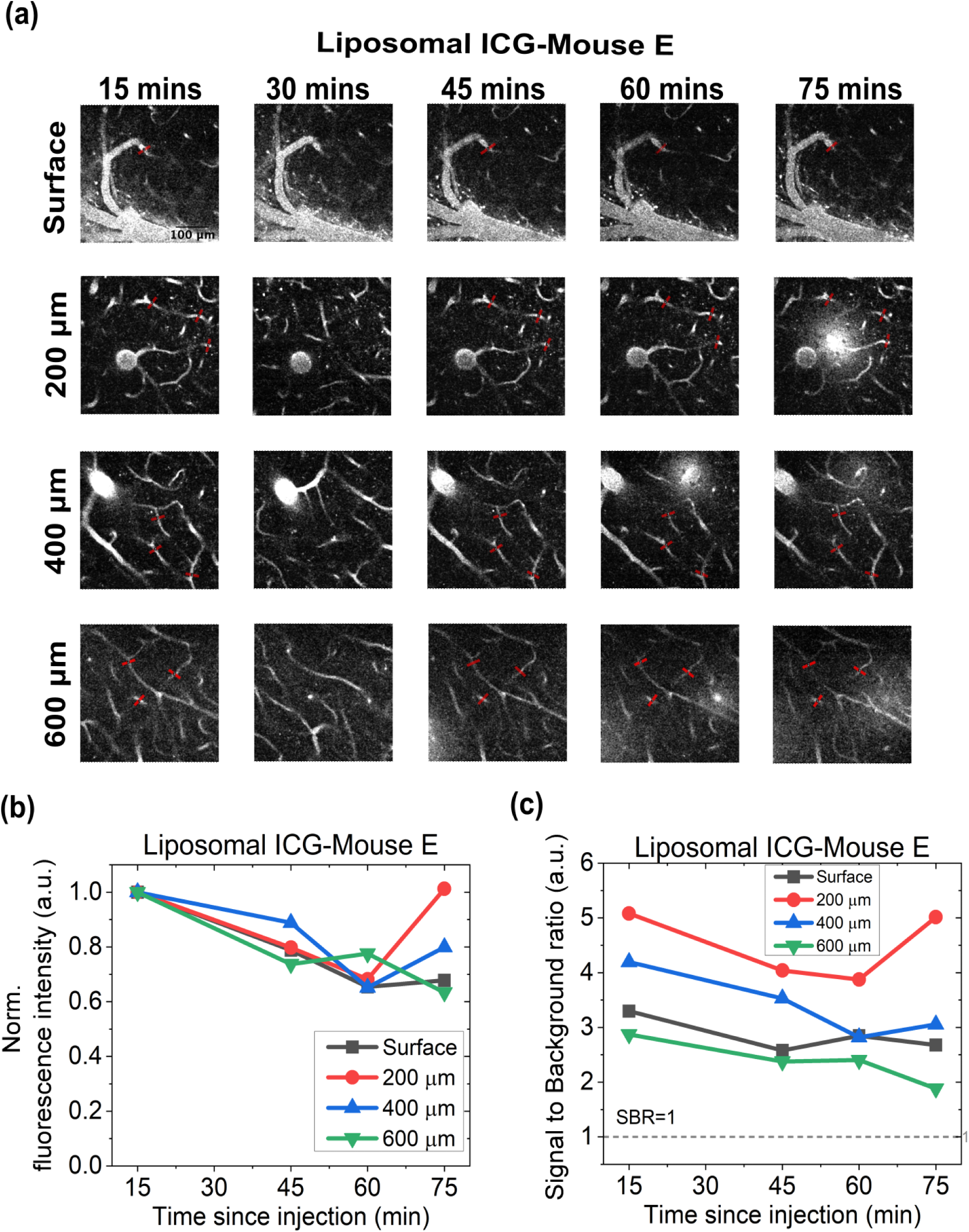
(a) *In vivo* imaging of blood vessels at different depths and time points for Mouse E using L-ICG. All images are displayed on the same intensity range. Scale bar represents 100 *µ*m. (b)Variation of overall mean fluorescence intensity for vessels at all depths. The overall mean fluorescence intensity at each time point is normalized to the value at the 15-minute time point. (c) Signal to background ratio (SBR) for different depths and time points. The SBR for a given depth at a specific time point was calculated by averaging the SBRs of 3 vessels at that depth. Only 1 vessel at the surface was chosen for analysis.

## 4 Discussion

Fluorescence imaging with ICG offers notable advantages, including low cost, accessibility, safety, and extensive availability of research. However, two-photon imaging of blood vessels labeled with ICG is limited by its relatively short clearance time. While a short imaging window of approximately 15 to 20 minutes may suffice for brief experiments, it is unsuitable for longer experiments that require deeper imaging or higher frame averaging. This time window is particularly limited, as it must accommodate positioning the rodent under the microscope and identifying a suitable cortical region for imaging.

A method to extend imaging duration with ICG by injecting a second retro-orbital dose to replenish the dye in the vessels, enabling deeper imaging. Although effective, this approach is not ideal, as re-injecting the dye once the mouse is positioned under the microscope is challenging. Furthermore, repeated injections may have undesirable effects on the rodent.

In this work, we propose liposomal encapsulation as a method to overcome the issue of rapid clearance of ICG from the rodent body. The mechanism of retention of ICG involves binding to the plasma proteins and consequent clearing by the liver or the kidney. We propose that the binding of the ICG particles to the plasma proteins can be prevented by encapsulating ICG in liposomal nanoparticles. In this way, the retention time for ICG in the rodent vascular network can be increased. Liposomal nanoparticles have been widely researched, are easily available, and are simple to synthesize. Further, the synthesis process adds negligible costs and is easily implementable through standard processes available in any wet lab.

As noted in Fig.S1, encapsulating ICG in liposomal carriers does not change the single-photon fluorescence intensity or absorbance spectra. Although we expected a similar trend for two-photon excitation, we wanted to verify this result. Figure S2 shows the normalized two-photon brightness curves for L-ICG and free ICG as a function of wavelength from 1250 nm to 1600 nm for different power levels. The curves show the range of wavelengths over which the fluorescent compounds can be efficiently excited via two-photon excitation. It is observed that the two-photon brightness spectra for L- and free ICG have similar trends. It is also concluded from these results that similar to free ICG, L-ICG can too be excited at 1300 nm. Thus, for all imaging purposes, we undertook two-photon excitation at 1300 nm.

Fig. 3(a) shows the results for the *in vivo* imaging procedure with free ICG. It is observed in the images that the vessels are clearly visible up until the 15 minute time point but they are not discernible from the 30 minute time point and beyond. This is attributed to the rapid clearance of free ICG from the mouse. At later time points, the vessel signal became indistinguishable from the background in the images, making it impossible to obtain any quantitative measurements of fluorescence in the vessels. Nonetheless, the signal from a single vessel was evaluated for the first two time points for the surface, 400 and 600 *µ*m depths. At 200 *µ*m, the signal from two vessels could be evaluated during the first two time points; however, only one vessel could be consistently tracked throughout the entire imaging session. Fig. 3(b) shows the variation of normalized fluorescence intensity as a function of time. The graphs in Fig. 3(b) show a significant drop in fluorescence intensities between 15 and 30 minutes, after which the vessels become indiscernible due to the clearance of ICG. Thus, it can be concluded that substantial clearance of ICG occurs within 15 to 20 minutes after injection. This is in agreement with what has been observed previously by Miller et al.^10^ There is one specific vessel at 200 *µ*m that was visible through the course of the entire imaging session. As seen in 3(b), the fluorescence intensity of this vessel decreases to 40% of its initial value at 15 minutes by the 30-minute mark and further declines to 20% by 45 minutes. The normalized fluroescence intensity then plateaus at approximately 20% and remains stable until the end of the imaging session at 75 minutes. Although this vessel remains visible for extended durations, it is an outlier, and any quantification of clearance from this vessel would clearly be an overestimation.

Fig. 4(a) shows the images obtained from *in vivo* experiments using L-ICG. It can be seen that at the same depth, the field of view is similar at all time points. Note that the imaging plane at the 30 minute time point was slightly shifted for depths of 200 *µ*m and beyond. For this reason, the images obtained at 30 min were excluded from the data analysis shown in Fig. 4(b) and (c). A key observation made from Fig. 4(a) is that the vessels are clearly distinguishable from the background for all time points and at all depths. This is impressive given that the same power levels are used to image all time points for a certain depth. Interestingly, at certain time points, such as the 75- minute mark at 200 *µ*m, vessel ablation is observed, possibly due to the high peak powers used for imaging. The issue of vessel ablation is beyond the scope of this study and will be addressed in detail in future research.

A quantitative understanding of the clearing can be obtained from Fig. 4(b). Figure 4(b) shows that for all depths the normalized fluorescence intensity decreases over the duration of the imaging exp‘eriment, but the decrease is gradual. Even at the 75-minute time point, the vessels were as bright as 60 80% of their initial brightness. The exception to this was the normalized intensity at a depth of 200 *µ*m. It was seen that at this depth the normalized fluorescence intensity increases compared to that at the start of the experiment. This is likely attributed to excess signal generated within the field of view, caused by vessel ablation near the vessels chosen for analysis.

Fig.4(b) shows the trend of the overall signal in the vessels over time but we also wanted to investigate affect of clearing on the signal to background (SBR) ratio. As seen in Fig. 4(c), the SBR gradually decreases over time. This can be attributed to the gradual loss in fluorescent signal from the vessels due to clearing of the dye. For all depths and at all time points, the SBR always stays well above 1. This signifies that the vessels are clearly distinguishable from the background in the field of view. Further, it is noticed that for this specific imaging experiment, the SBR for the surface vessels is lower than that for the deeper depths. This is not an expected result and the following paragraph describes our understanding of why that happens.

The signal-to-background ratio (SBR) is influenced by two key factors: (i) the mean signal intensity from the vessels and (ii) the mean background intensity in the vicinity of the vessel. Consequently, SBR is highly sensitive to variations in the background. From the analysis, it was observed that the same vessel at different time points could exhibit significant variance in the mean background. This may result from artifacts near the vessel, causing an unexpectedly high background, or from extremely low background levels, leading to an unusually high SBR. Therefore, while generating line profiles, careful attention was paid to avoid regions on the vessels that might encounter either of these issues. It was difficult to completely avoid this and so the absolute values for the SBRs at different depths can not be compared accurately. Since our focus is on assessing dye clearance, we are primarily concerned with the trend of SBR over time at various depths rather than their exact values.

By comparing the images in Figures 3 and 4, it can be seen that liposomal encapsulation leads to increased retention of ICG within the blood vessels. In contrast to the observed clearance time of 20 minutes with free ICG, encapsulation allows ICG to remain in the vessels for up to 75 minutes. It is reasonable to believe that the dye would have remained in the vascular network for over 75 minutes; however, this was not investigated in the current study. A longer imaging window would enable a smoother experimental protocol and facilitate experiments requiring extended durations, such as deep vascular imaging.

## 5 Conclusion

Indocyanine Green is an FDA-approved, accessible vascular label for two-photon imaging. Its rapid clearance from the rodent body serves as one of the limiting factors for *in vivo* two-photon imaging. We propose a safe and inexpensive technique to improve the retention of ICG that involves the encapsulation of ICG particles in liposomal nanoparticles. We compare the clearance time for non-encapsulated or free ICG and L-ICG for *in vivo* imaging and note that encapsulation improves the clearance time for ICG from 20 minutes to atleast 75 minutes. This work is an attempt towards improving the methods for two-photon microscopy with ICG. Given the wide prevalence of ICG in clinical settings, these improved methods could be vital towards translating two-photon microscopy to the clinic.

**Alankrit Tomar** is a Senior Systems Engineer at the University of Texas at Austin. He received his BTech. degree in Engineering Physics from Delhi Technological University in 2019, and his MS and PhD degree in Electrical and Computer Engineering from University of Texas at Austin. His PhD. work focuses on improving methods for fluorescent dyes for two-photon microscopy of cerebrovascular structures. He has been a member of SPIE and Optical Society of America (OSA, now Optica).

**Noah Stern** is a PhD Candidate in the Department of Biomedical Engineering at the University of Texas at Austin. He is a member of the Diverse Engineering Applications Lab (DEAL) under the supervision of Dr. Tyrone Porter. His research focuses on novel theragnostic nanoparticles as contrast agents for fluorescence and photoacoustic imaging as well as photothermal therapy.

Biographies and photographs of the other authors are not available.

