## Supplementary material for "Improving retention time of Indocyanine Green for *in vivo* two-photon microscopy using liposomal encapsulation"

### Synthesis of ICG-encapsulated liposomal nanoparticles:

Promising results with multiple different imaging modalities and use cases suggest that multi-photon imaging utilizing L-ICG may be equally as successful in prolonging imaging time. To achieve this, an optimized procedure for producing L-ICG has been developed that maximizes indocyanine green encapsulation, improves stability, and maintains a sufficient PEG coating to increase circulation time. The lipids chosen for creating this formulation of L-ICG are Distearoylphosphatidylcholine (DSPC), cholesterol, and 1,2-distearoyl-sn-glycero-3-phosphoethanolamine-N-[amino(polyethylene glycol)-2000] (DSPE-PEG2K). DSPC is the major phospholipid component of the liposome chosen for its relatively high transition temperature (55.6°C), ease of handling, low cost, and abundance in the literature. A higher transition temperature should limit leakage of ICG once L-ICG are administered *in vivo*. Cholesterol is included at a relatively high ratio to improve the stability and fluidity of the liposome at higher temperatures. DSPE-PEG2K is included at the above ratio to enable viable PEG "brush" formation. A schematic of the formulation process is shown in Figure 1 of the manuscript.

Briefly, the process can be broken down in two phases: thin-film hydration and liposome formation. Thin-film hydration is a well-characterized and relatively standardized technique commonly employed for liposome synthesis.[1] In this process, lipids dissolved in a volatile organic solvent like chloroform are added to a vial or round-bottom flask in a pre-defined and optimized ratio. For DSPC, cholesterol, and DSPE-PEG2K, this ratio was found to be optimal at 7:3:1. Changes to cholesterol percentage may disrupt the fluidity of the liposome, too high of a PEG concentration can inhibit particle formation, while too low of a PEG concentration can limit the efficacy of the PEG "brush." Once in the flask, lipids are gently agitated under a stream of inert gas or placed in a rotary evaporator to remove the organic solvent. If successful, following evaporation a thin cloudy film of lipid will be left along the wall of the flask. Next, the thin-film is hydrated with a solution containing the desired cargo. As the solution is added, the film will swell and spontaneously peel off from the wall forming a polydisperse collection of spherical multilamellar vesicles. As they swell and detach from the walls of the flask, they will encapsulate portions of the hydrating solution. In this case, the hydrating solution contains ICG at a defined concentration (~1.0 mg/ml). The concentration is key, as a higher initial concentration means more ICG will end up inside the forming vesicles. However, going too high can inhibit vesicle formation or begin to instigate the formation of ICG J-Aggregates (ICG-JA).[2,3] ICG-JA are a more stable form of ICG that can form at high concentrations and temperatures. They have higher levels of absorbance, but significantly lower fluorescence intensity than ICG.[4] After hydration, the resulting solution is heated near the transition temperature of DSPC while cyclically vortexing and sonicating to ensure all lipid is detached and to increase encapsulation efficiency. The result is a solution of multilamellar vesicles containing ICG in a solution also containing free floating ICG.

The second phase of the L-ICG synthesis is liposome formation. The first step of this phase is to extrude the solution of multilamellar vesicles down in to small unilamellar vesicles or liposomes. Extrusion is another well-established process by which solutions of particles are repeatedly pushed through polycarbonate membranes with defined pore sizes.[5] As larger particles are pressed up against the membrane they are forced through the smaller pores cutting them down to size and removing extra lipid layers. The more cycles through the membrane, the closer the average particle diameter gets to the pore size and the lower the polydispersity index (PDI). For this application, a pore size of 100 nm was chosen and liposomes were extruded at least 11 times. Following extrusion, the particles are dialyzed against ultrapure water at 4°C for 48 hours with the water being changed every 16 to 24 hours. Dialysis removes any unencapsulated ICG from the solution so the only ICG signal will come from molecules encapsulated within the liposomes. Dialyzing at 4°C is also essential to protect the stability of the liposomes and again limit any chance that J-aggregation will begin. The final step is to filter the liposome solution through 0.22μm sterile filters to remove any contaminants or larger aggregates left over from extrusion or aggregation. At this point the particles are ready for characterization. If they are to be used for *in vivo* experimentation an additional step involving centrifugation using Amicon® filters is performed. The pellet is resuspended to the proper concentration for injection using sterile saline or PBS.

### Characterization of ICG-encapsulated liposomal nanoparticles:

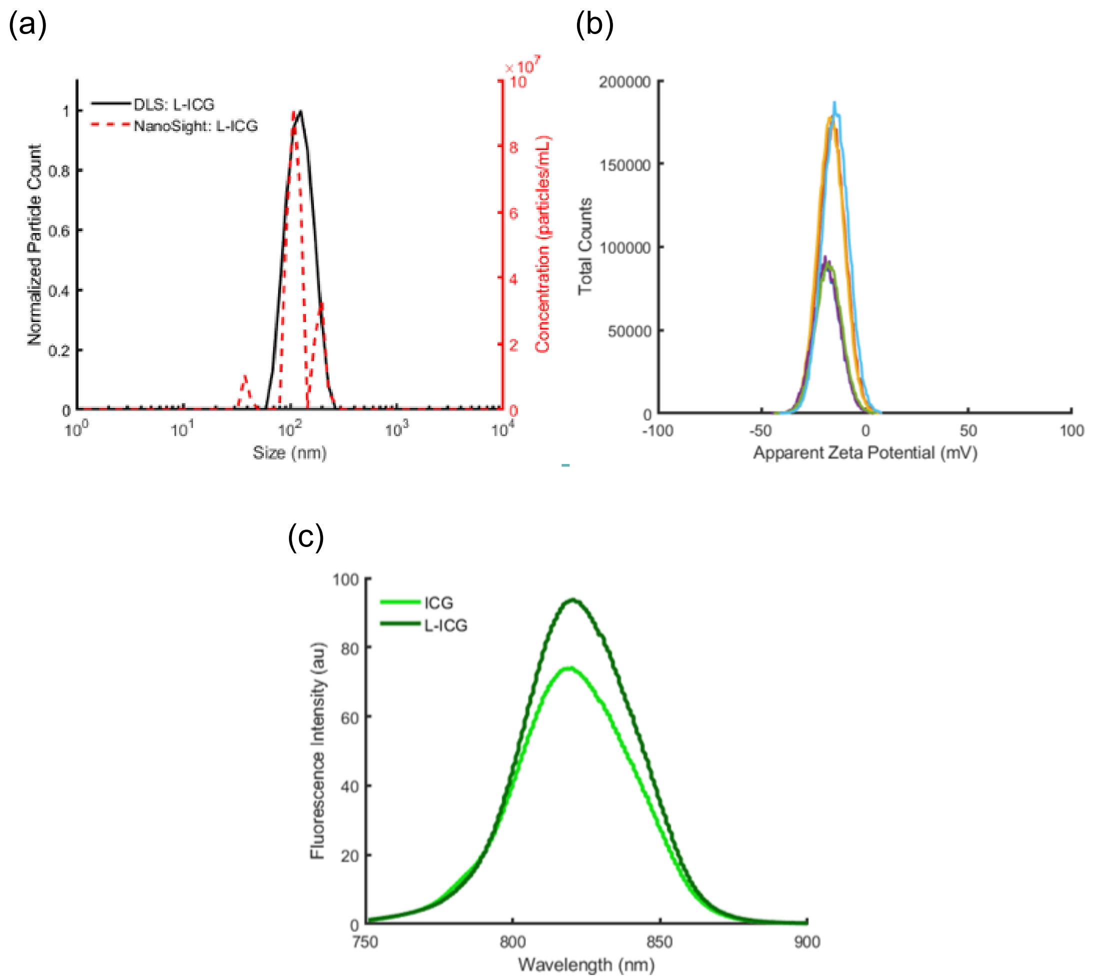


**Figure S1: Liposomal-ICG Characterization**. Before imaging, (a) the size, (b) zeta potential, (c) single photon fluorescence intensity are measured. L-ICG have acceptable negative zeta potentials of sufficient magnitude to prevent aggregation. L-ICG have similar fluorescence properties to ICG while producing a stronger intensity.

To fully characterize the particles, their final encapsulation efficiency, fluorescence intensity, absorbance, size, and zeta potential are determined each production cycle to ensure proper synthesis and maintain consistency across batches. Various characterization results are included in Figure S1 . Encapsulation efficiency describes the amount of ICG that remains in solution relative to the initial concentration used to hydrate the lipids during formulation. This is typically performed by calculating the ratio of absorbance intensities at peak wavelengths. However, ICG has the peculiar quality of having concentration dependent absorbance peaks that correspond to dimerization of ICG.[6] To overcome this quality, a one-to-one dilution of the particles to 100% ethanol is created. In ethanol, all ICG dimers are dissolved into monomers so the absorbance signal will be consistent each time. Absorbance measurements of the ethanol containing solution are compared to the same measurements taken from the initial ICG hydration solution to calculate an encapsulation efficiency. For L-ICG using this method, the encapsulation efficiency comes to 21.41 ± 5.28%. This same technique is used to calculate the concentration of ICG in the liposomes following formation. Instead of comparing to the initial hydration solution, the absorbance values are compared to a calibration curve built using known concentrations of ICG in ethanol. Fluorescence intensity and absorbance measurements are taken to ensure there are no significant changes compared to free ICG. Other key changes to observe would be a shift in peak absorbance wavelength or a significant drop in fluorescence intensity indicative of J-aggregate formation (Figure S1c). The liposomal shell does not appear to alter L-ICG fluorescence wavelengths from ICG, but the high concentration of ICG within each particle does appear to increase fluorescence intensity slightly. Size is determined using Dynamic Light Scattering (DLS). Pegylated nanoparticles between the sizes of 50 and 200nm will typically circulate the longest and most efficiently as they are too large for immediate kidney clearance, too small for immediate uptake in the liver, and temporarily hidden from immune cells of the reticuloendothelial system. Typically, it is also best to aim for a PDI of 0.35 or lower. A lower PDI indicates a more uniform sample that is likely to be more stable over time.[7] For L-ICG using this method, average sizes come to 124.04 ± 12.39nm with a PDI of 0.312 ± 0.021 (Figure S1a). NanoSight tracking was also employed to characterize particle size and concentration. Average size measurements align well with DLS results, and particle concentrations appear to be on the order of $10^8$ particles per milliliter (Figure S1a). Finally, zeta potential is a measure of the potential difference between the particle surface and its background solution.[8] Generally, it can be thought of as an indicator of particle stability over time and potential toxicity. Zeta potentials between -10mV and +10mV typically indicate neutral solutions which may aggregate over time. Higher magnitudes near 30mV are generally more electrostatically stable as they are unlikely to aggregate, but high magnitude positive potentials are often associated with heightened risk for toxicity. Moving to a more negative/neutral potential without dropping below a 10mV magnitude will ideally prevent long-term aggregation and improve stability[9] while maintaining low toxicity.[10] For L-ICG using this method, average zeta potentials come to -16.47 ± 1.43mV (Figure S1b).

### Two-photon relative brightness spectra for free ICG and liposomal-encapsulated ICG

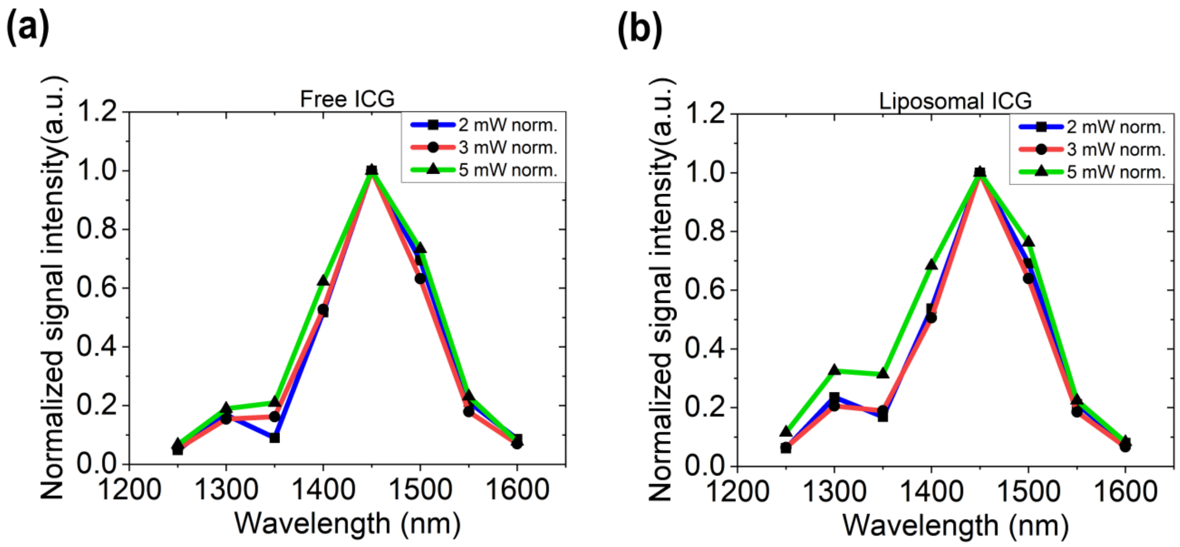


**Figure S2:** Normalized two-photon excitation spectra for (a) free ICG and (b) L-ICG using different power levels

### Imaging results using L-ICG in two other animals:

(a)

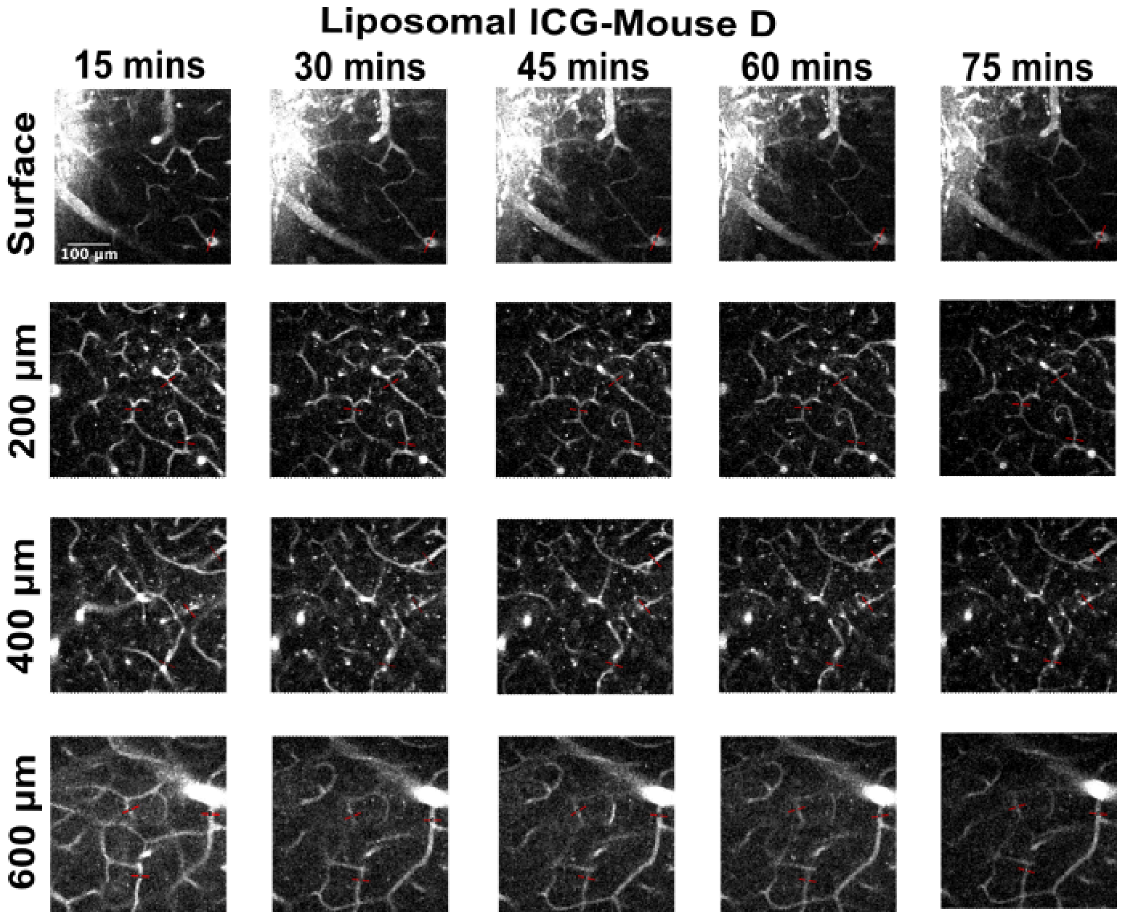


(b)

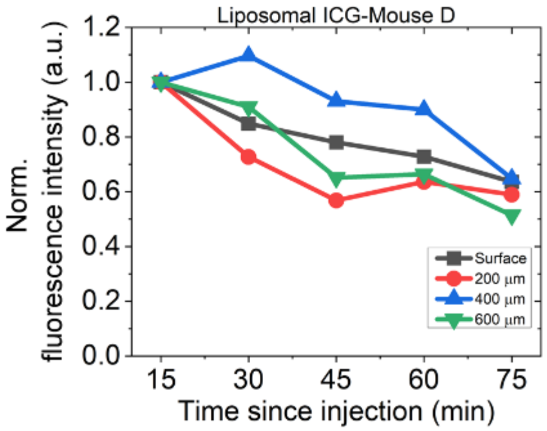


**Figure S3:** (a) *In vivo* imaging of blood vessels at different depths and time points for Mouse D using L-ICG. All images are displayed on the same intensity range. Scale bar represents 100 µm.

(b)Variation of overall mean fluorescence intensity for vessels at all depths. The overall mean fluorescence intensity at each time point is normalized to the value at the 15-minute time point.

(a)

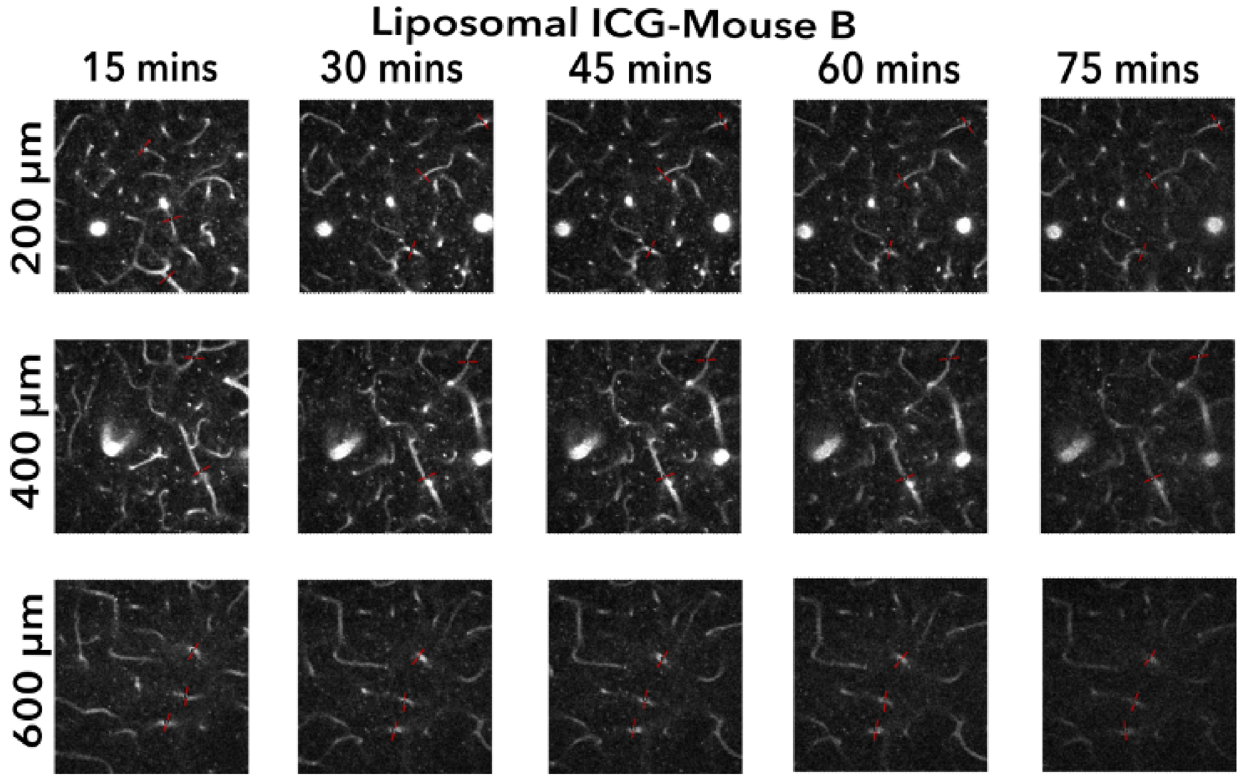


(b)

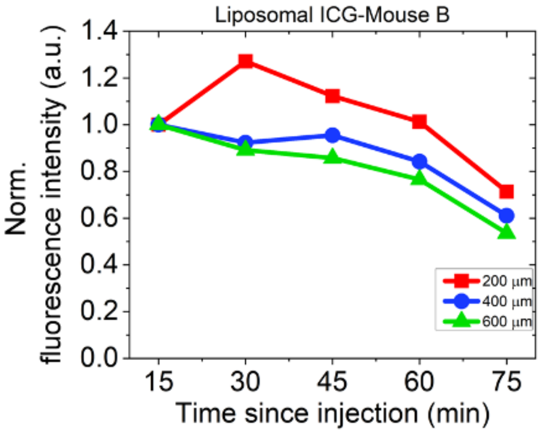


**Figure S4:** (a) *In vivo* imaging of blood vessels at different depths and time points for Mouse B using L-ICG. All images are displayed on the same intensity range. Field of view is the same as other vascular images (b)Variation of overall mean fluorescence intensity for vessels at all depths. The overall mean fluorescence intensity at each time point is normalized to the value at the 15-minute time point.
